# Single-cell heterogeneity in interferon induction potential is heritable and governed by variation in cell state

**DOI:** 10.64898/2025.12.09.693293

**Authors:** Elizabeth A. Thayer, Gabriela Shipman, Joel Rivera-Cardona, Tarun Mahajan, Qi Wen Teo, J. Sebastian Paez, Joseph Lederer, Jie Chen, Nicholas C. Wu, Sergei Maslov, Christopher B. Brooke

## Abstract

Type I and III interferons (IFNs) are among the first lines of defense against viral infection, yet they are generally only produced by a tiny fraction of infected cells. Here, we show that cellular heterogeneity in IFN induction potential upon treatment with immunostimulatory RNA is not due to variability in sensing of stimuli but instead is shaped by heterogeneity in tonic cell signaling state. Using complementary single-cell approaches, we found that baseline variation in the c-Jun N-terminal kinase (JNK) and activator protein (AP)-1 transcription factor families correlated with *IFNL1* expression predisposition. We further show that drug-based inhibition of JNK signaling virtually eliminates the innate antiviral response to immunostimulatory RNA. Finally, we show that single cell heterogeneity in IFN induction potential is heritable and stably maintained over numerous generations. Together, our study emphasizes the influence of intrinsic variability in cell state on innate immune regulation and IFN induction heterogeneity.

## INTRODUCTION

The induction of interferon (IFN) is a critical determinant of infection outcome for a wide variety of viruses^1–5^. Respiratory epithelial cells express both type I and III IFNs (IFNα/β and IFNλ, respectively) as part of their first line of defense against the replication and spread of viruses like influenza A virus (IAV). While most cells in the body can respond to type I IFN, only epithelial cells and a subset of immune cells express the correct receptor for responding to type III IFN^3,6^. Due to this tissue specificity in responsiveness, IFNλ is thought to be less inflammatory while still inducing the expression of IFN-stimulated genes (ISGs) that are important for cellular antiviral defense^1,6^.

Retinoic acid-inducible gene I (RIG-I) and melanoma differentiation-associated protein 5 (MDA5) are cytosolic pathogen recognition receptors that recognize viral and synthetic foreign RNAs with distinct molecular features^7–9^. RIG-I and MDA5 share similar downstream signaling pathways, as they activate mitochondrial antiviral-signaling protein (MAVS) and ultimately lead to the nuclear translocation of interferon regulatory factor 3 (IRF3) and the initiation of IFN transcription^6,8^. We and others have demonstrated that only a small percentage of infected cells typically express IFNs, independent of virus type^10–15^. The specific factors contributing to heterogeneity in IFN-induction potential remain poorly understood.

The existence of substantial phenotypic heterogeneity within cellular populations, even for cells of a given “type”, is a well-established phenomenon. Many studies have investigated how cell-intrinsic factors, such as cell cycle stage, mitogen-activated protein kinase (MAPK) signaling state, and patterns of chromatin accessibility, influence variability in response to a stimulus or other cellular phenotypes^16–21^. Intrinsic 2’-5’-oligoadenylate synthase-like protein (OASL) expression has been recently identified as a critical factor governing cellular heterogeneity in IFN induction potential during IAV infection, but it is not clear whether this effect is IAV-specific^22^.

Here, we investigated the sources of cellular heterogeneity in the ability of A549 cells to produce IFN in response to the commonly used RIG-I agonist, polyinosinic:polycytidylic acid (pIC). We used single-cell RNA sequencing and pseudotime analysis to identify transcriptional determinants of IFN induction potential. Additionally, we found that single-cell clonal populations isolated from a single parental population exhibited massive differences in IFN-induction phenotypes that were maintained over numerous generations. This heritable phenotypic variation correlated in part with basal transcriptional and epigenetic profiles. Using multiple approaches, we identified cellular variation in (a) basal expression of ISGs, and (b) MAPK-activator protein (AP)-1 signaling activity as key predictors of the ability of individual cells to mount a robust innate immune response to foreign RNA.

## RESULTS

### Following pIC treatment, most cells exhibit IRF3 nuclear translocation but do not transcribe IFNλ

Despite the critical importance of the IFN response for host defense, we and others have demonstrated that only a small fraction of cells are generally capable of expressing IFN following viral infection^10–13,23–25^. Viral infection dynamics are highly heterogeneous at the single-cell level, and it is unclear how much of the observed variation in IFN induction is due to cellular heterogeneity in viral processes^14,26,27^.

To examine single-cell heterogeneity in IFN induction without the confounding effects of viral heterogeneity, we treated the human lung adenocarcinoma cell line A549 with increasing concentrations of pIC and quantified *IFNL1* transcription within single cells using hybridization chain reaction^28^ fluorescence *in situ* hybridization coupled with flow cytometry (HCR-Flow)^29^. Even after 24 hours post-treatment with the highest dose tested, fewer than 20% of cells expressed detectable levels of *IFNL1* (**Fig. 1A**). Thus, most cells are incapable of expressing IFN in response to pIC, regardless of dosage or timing.

**Figure 1.**
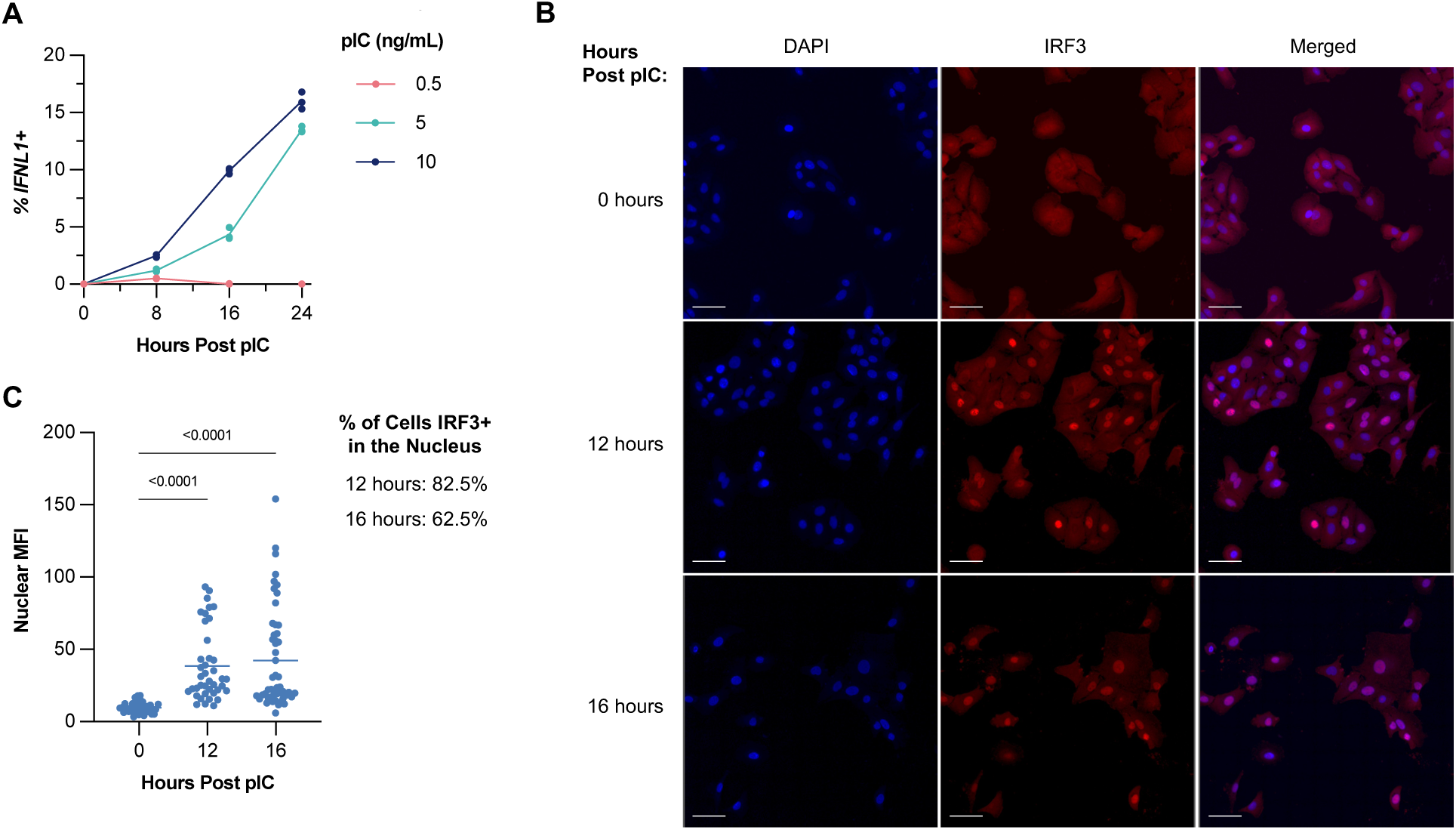
IFNL1 is rarely expressed in A549 cells after pIC stimulation. **(A)** Percent of cells positive for IFNL1 transcripts at 0-, 8-, 16-, and 24-hours post-pIC treatment with 0.5, 5, and 10 ng/mL concentrations. **(B)** Representative images of A549 cells stimulated with pIC with the nucleus and IRF3 stained. White bars indicate 50 μm. Brightness enhanced evenly for publication. **(C)** Quantification of raw fluorescence of IRF3 in the nucleus using Imaris. Significance calculated using Dunnett’s multiple comparisons. Percent of cells IRF3+ in the nucleus determined by the number of cells in the stimulated populations with a higher MFI than the highest untreated cell.

Activation of pattern recognition receptors like RIG-I or toll-like receptor 3 (TLR3) typically triggers a signaling cascade that results in the phosphorylation and nuclear translocation of IRF3, the primary transcription factor responsible for driving type I and type III IFN induction^6^. If the low frequency of *IFNL1* induction we observed following pIC stimulation was due to a failure to deliver or detect pIC, then we would expect that IRF3 translocation would occur at similarly low frequencies. We treated A549 cells with pIC and measured the percentage of cells in which IRF3 translocated to the nucleus using confocal fluorescent microscopy. We observed IRF3 nuclear translocation in most pIC-treated cells at 12- and 16-hours post-treatment (**Fig. 1B,C**). These results suggest that the low frequency of *IFNL1* induction cannot simply be explained by defects in sensing of pIC or canonical IRF3 signaling.

### Gene expression profile prior to treatment is predictive of *IFNL1* induction following pIC treatment

Pseudotime analysis has previously been used to identify genes whose baseline expression at the single-cell level correlates with the likelihood that a given cell expresses *IFNL1* in response to IAV infection^22,30^. Since active IRF3 signaling alone is insufficient for robust *IFNL1* expression after pIC stimulation, we sought to use this approach to identify other crucial contributors to the *IFNL1* response following pIC treatment. We isolated A549 cells at 0-, 4-, 8-, 12-, and 16-hours post-pIC treatment for single-cell RNA sequencing (scRNA-seq) to then infer the velocity of individual cells moving through the UMAP dimensions (**Fig. 2A**). This approach identified five terminal states at 16 hours, two of which were *IFNL1*+ (**Fig. 2B**).

**Figure 2.**
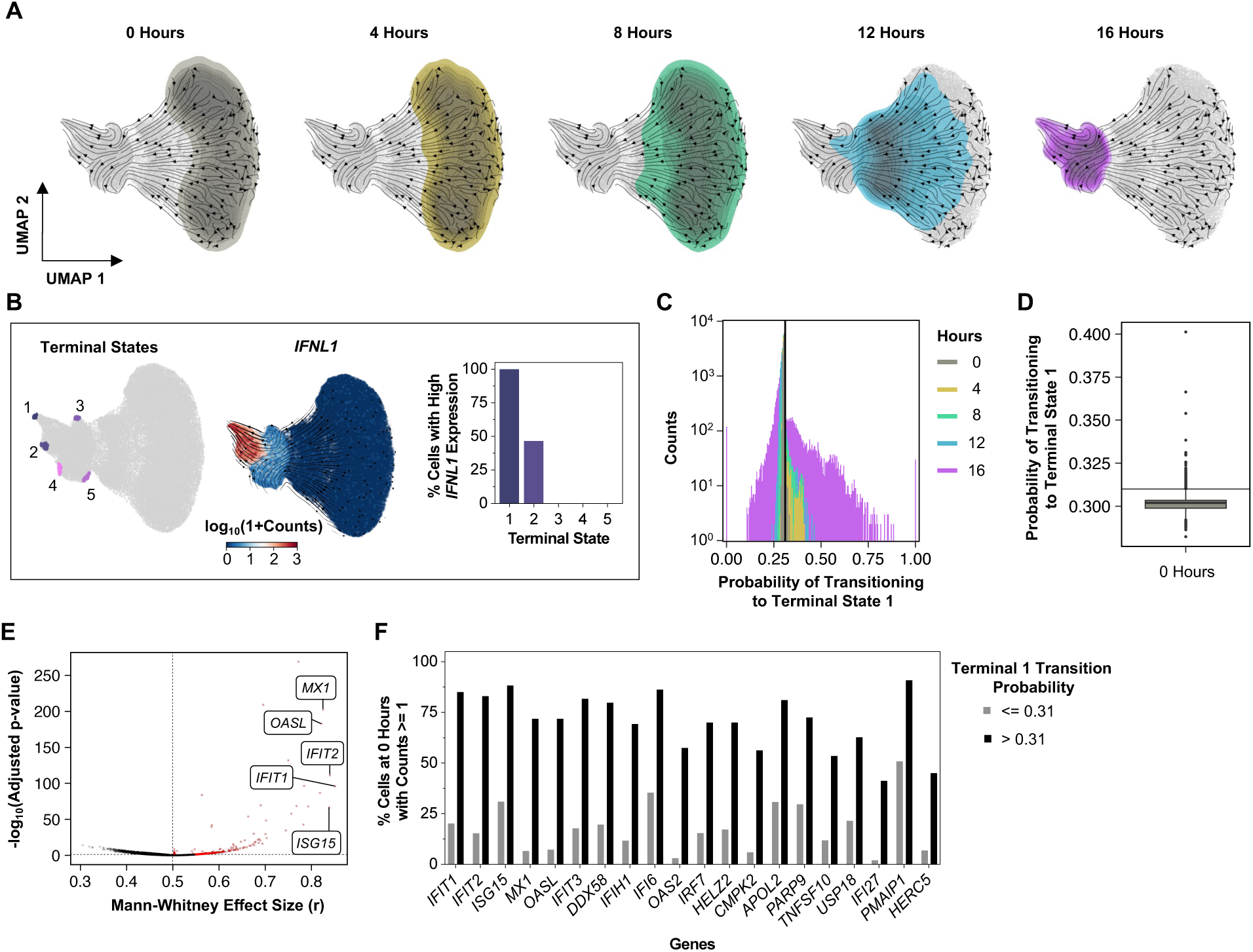
Charting the single-cell transcriptional response to pIC using RNA velocity. **(A)** UMAP dimensional reduction of the cells from each time point aggregated together, then colored by time point – time indicating sample collection after 5 ng/mL pIC treatment. **(B)** UMAP dimensional reduction of all cells from every time point. (Left) Terminal states were calculated using temporal-noSpliceVelo and used to determine the probability that each cell in the dataset would end up in those terminal states. (Middle) UMAP dimensional reduction of all cells colored by log10(IFNL1 Counts). (Right) Table showing the percent of cells with IFNL1 counts >= 5 within each terminal state (probability = 1). **(C)** Distribution of the Terminal 1 transition probability of cells in each collection time point. Line at 0.31 indicating the mean transition probability among the full dataset, ultimately being the threshold for high-probability cells. **(D)** Distribution of the terminal 1 transition probability per cell in the 0-hour collection time point. Line at 0.31, the previously established as the threshold for high-probability cells. **(E)** Visualization of the genes significantly more expressed in the high-probability cells at 0 hours. Points in black are the unfiltered genes, and points in red are filtered genes - log10(adjusted p-value) > 1.3 and Mann-Whitney effect size > 0.5. **(F)** Percent of cells at 0 hours expressing the top 20 genes from the Mann-Whitney analysis, separated by the cells’ Terminal 1 transition probabilities.

We next inferred the probability of each individual cell ending up in terminal state 1, the only terminal state where cells were 100% *IFNL1*+ (**Fig. 2B**). The probabilities are not normally distributed and become increasingly bimodal with time, making a correlation analysis between the probabilities and the gene expression of each cell suboptimal.

Instead, we set a threshold for high- and low-probability cells, where “high” is classified as having greater than the mean probability across all time points (**Fig. 2C**). This threshold, when plotted on the 0-hour distribution alone, showed that the high-probability cells were outliers, which was consistent with the rarity of *IFNL1*+ cells (**Fig. 2D**). We used the top 4,000 most variable genes across the time points to conduct a Mann-Whitney test to identify genes that are more likely to be expressed in the high-probability cells at 0 hours than the low-probability cells.

This analysis identified 301 genes expressed at the 0-hour time point with an adjusted p-value lower than 0.05 and an effect size greater than 0.5, indicating they are more likely to be expressed in high-probability cells (**Fig. 2E**). Upon further analysis, we could visualize the result of the test when comparing the percentage of high- or low-probability cells at 0 hours expressing the top 20 genes. All but two of the top genes were expressed in most of the high-probability cells at 0 hours, with all the genes expressed in a greater percentage of high-probability cells (**Fig. 2F**). Additionally, all of the top 20 genes have been reported as ISGs, consistent with the emerging idea that intrinsic ISG expression and/or tonic IFN signaling governs IFN induction potential^22,31^.

### Single-cell clones exhibit substantial heritable variation in IFN induction potential

Our results indicated that pre-existing heterogeneity within cell populations underlies variability in IFN-induction potential. We next asked whether this variability in IFN-induction potential was transient or if it could be maintained over multiple generations. To distinguish between these possibilities, we sorted single cells from a population of A549 cells and expanded them into 11 individual clonal lines, denoted ET1-ET11.

We treated the parental cell population and the individual clonal lines with 0.5 ng/mL pIC and quantified *IFNL1* induction at 16 hours post-treatment. *IFNL1* transcript levels varied nearly 1000-fold between the individual clonal lines (**Fig. 3A**). Thus, the individual clonal lines exhibited substantial variation in pIC responsiveness that was maintained through the numerous rounds of cell division required to expand them.

**Figure 3.**
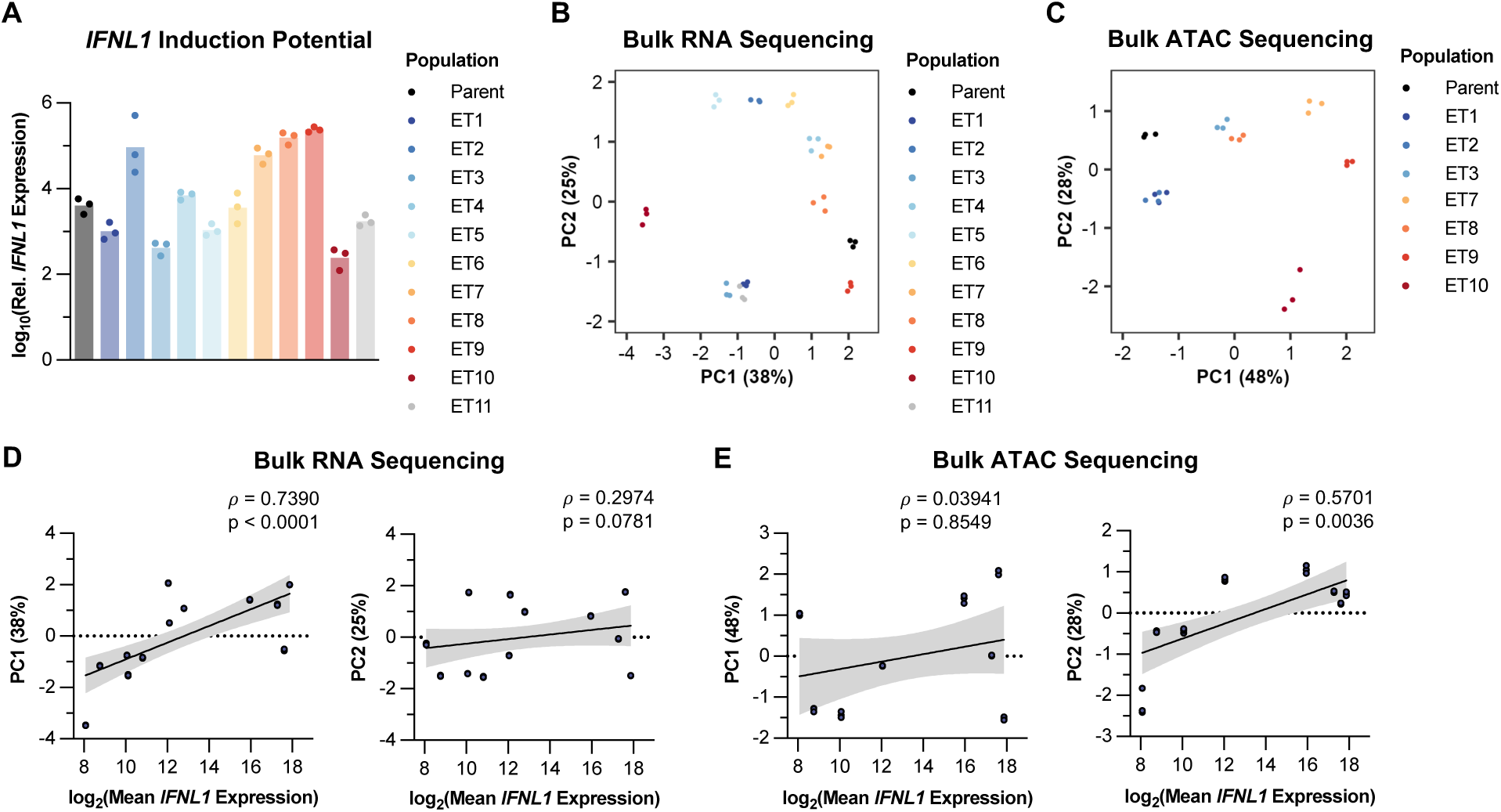
Sibling clonal cell lines exhibit substantial heritable variability in IFN induction phenotypes. **(A)** IFNL1 expression 16 hours post 0.5 ng/mL pIC treatment among the cell lines was quantified using RT-qPCR. Relative log10(ddCt) fold changes to their ACTB expression and the mean IFNL1 level from the mock of each respective population. One-way ANOVA analysis p < 0.0001. **(B)** PCA plot comparing the transcriptional profiles of each bulk population. **(C)** PCA plot comparing the chromatin accessibility profiles of each bulk population. **(D)** Correlation between the PC1 (left) and PC2 (right) components of the bulk RNA sequencing data and the log2(Mean IFNL1 Expression) of each of the clonal lines. Trendline and 95% confidence interval generated with simple linear regression. ρ and p-value calculated based on Spearman’s correlation. **(E)** Correlation between the PC1 (left) and PC2 (right) components of the bulk ATAC sequencing data and the log2(Mean IFNL1 Expression) of each of the clonal lines. Trendline and 95% confidence interval generated with simple linear regression. ρ and p-value calculated based on Spearman’s correlation.

In parallel, we compared the basal transcriptional profiles of the individual clonal lines in the absence of treatment using bulk RNA sequencing and observed that most of the clonal lines diverged substantially from the parental population (**Fig. 3B**). We compared the PC1 component with *IFNL1* induction phenotypes and found a strong correlation between basal transcriptional state and *IFNL1* induction potential (**Fig. 3D**).

The apparent heritability of heterogeneity in IFN induction phenotypes over numerous generations led us to ask whether this phenotypic variability correlated with any epigenetic signatures. We compared patterns of chromosomal accessibility across the clonal lines using bulk Assay for Transposase-Accessible Chromatin with sequencing (ATAC-seq). We found that global patterns of chromatin accessibility were highly distinct across the clonal lines (**Fig. 3C**). Like what we observed for patterns of transcriptional similarity, ET10 (low *IFNL1* induction) was separated from the group and clearly distinct from ET9 (high *IFNL1* induction). We then asked if variation in the lines’ accessibility profiles related to their *IFNL1* responses (**Fig. 3E**). While we did not find a significant correlation between the chromosomal accessibility PC1 and the magnitude of the *IFNL1* response, there was a significant correlation when using the PC2 coordinates (**Fig. 3E**). This suggests that, although the main factor contributing to the heterogeneity in the ATAC data may not be linked to *IFNL1* predisposition, there is significant connection between chromatin accessibility patterns and *IFNL1* expression potential.

### Variation in basal expression of RIG-I pathway components does not explain the heterogeneity in *IFNL1* induction phenotypes

We next sought to determine whether the basal expression patterns of individual transcripts in the clonal populations could also explain heterogeneity in the IFN response to pIC, as we observed in our longitudinal scRNA-seq dataset. We used a generalized linear model, where the x-axis values were the mean log2 fold changes in *IFNL1* expression of the clonal lines following pIC treatment, and the y-axis values represented the relative expression levels of individual transcripts in the same cell lines in the absence of treatment.

We first examined the transcripts that encode proteins known to be centrally involved in the RIG-I-based RNA-sensing pathway: *DDX58*, *MAVS*, *TBK1*, and *IRF3*^8,32^ (**Fig. 4A,B**). We focused specifically on RIG-I because knocking down other RNA sensors, MDA5 (*IFIH1*) and *TLR3*, did not reduce *IFNL1* expression after pIC stimulation as we saw with *DDX58* knockdown, suggesting that the response to pIC in this system is RIG-I-dependent (**Fig. 4A**). While *DDX58, MAVS,* and *TBK1* transcripts did exhibit significant positive correlations with *IFNL1*-induction potential, they only varied in expression levels by 2- to 4-fold across the clonal lines. The expression of the main transcription factor thought to be involved in initiating *IFNL1* expression, *IRF3*, was not significantly correlated with *IFNL1* across the cell lines. Overall, these weak trends appear insufficient to explain the nearly 1000-fold range in IFN expression levels we observed across the clonal lines.

**Figure 4.**
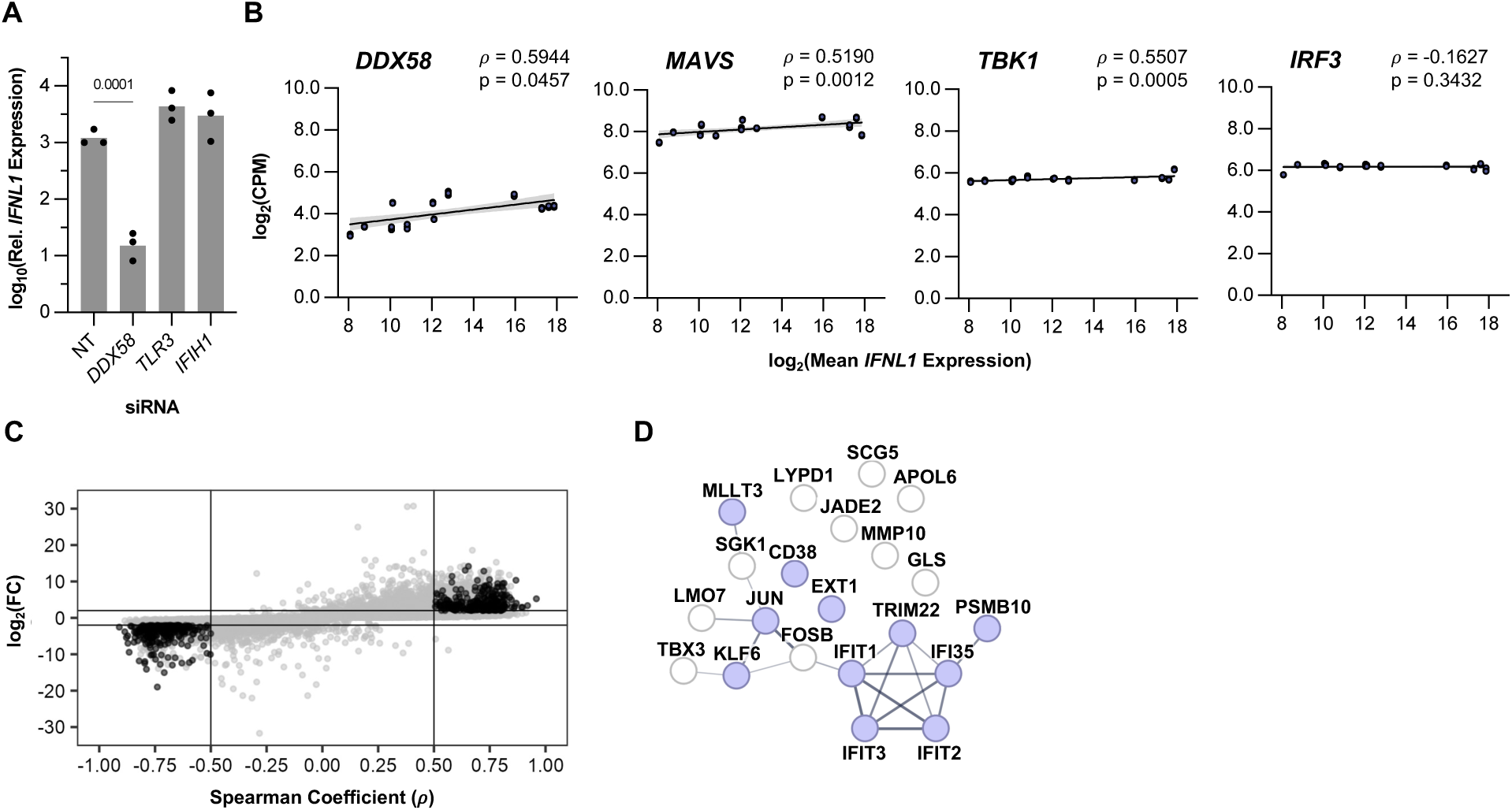
Identification of genes showing a correlation with IFNL1-induction potential. **(A)** IFNL1 expression 16 hours post 0.5 ng/mL pIC treatment after 48-hour siRNA treatment was quantified using RT-qPCR. Relative log10(ddCt) fold changes to their ACTB expression and the mean IFNL1 level from the mock of the non-targeting (NT) control. Significance determined by Dunnett’s multiple comparisons test. **(B)** Gene trajectory analysis correlating pIC-induced IFNL1 expression with basal gene expression. Trendline and 95% confidence interval generated with simple linear regression. ρ and p-value generated using Spearman’s correlation. **(C)** Plot of the genes following the generalized linear model (grey), with the genes passing set thresholds of |ρ| > 0.5, -log10(FDR) > 3, |log2(FC)| > 2, and log2(CPM) > 0.1 (black). **(D)** STRING analysis of the top genes found in the clonal bulk RNA sequencing and the single-cell data (PPI enrichment p < 0.0001). The thickness of the edges indicates the strength of data support for the function and physical protein associations (interaction score > 0.4). Nodes colored purple are annotated to be involved in immune system processes.

To identify gene products outside of the canonical RIG-I/MAVS pathway that could modulate IFN induction potential, we identified all transcripts that exhibited both (a) a linear trend (|ρ| > 0.5) between expression level and *IFNL1* induction, and (b) a wide range of expression (|log2FC| > 2) across the clonal lines (**Fig. 4C**). Over 500 genes passed these filtering criteria, so to narrow down the list we examined the common genes present in both the clone and longitudinal scRNAseq datasets. There were 21 genes in both lists, with significantly more interactions^33^ than expected (PPI enrichment p < 0.0001), suggesting a significant biological connection (**Fig. 4D**). Indeed, there are known ISGs in the list, *TRIM22*, *IFIT1*, *IFIT3*, *IFIT2*, and *IFI35*^34–36^, consistent with the observation that basal ISG expression is associated with enhanced IFN induction potential^22,31^. Notably, two components of the AP-1 complex, *JUN* and *FOSB* were also on this list. While cJun is known to bind to the IFNB enhanceosome^37,38^, AP-1 signaling has not been implicated in modulating type III IFN expression. These data raised the possibility of AP-1 signaling as another source of heterogeneity in IFN-induction potential.

### MAPK pathway signaling status modulates *IFNL1* induction potential

AP-1 activity is heavily influenced by upstream MAPK signaling and is potently activated by tumor necrosis factor (TNF)^39–42^. To examine the role of AP-1 activity in IFN induction heterogeneity in our system, we first asked how activation of MAPK signaling via TNF pretreatment affected IFN induction by pIC in both the parental A549 cell population and the ET9 clonal line that exhibited a high IFN induction phenotype. However, we wanted to focus only on TNF’s potential role in *IFNL1* induction, so we used ruxolitinib^43^ pre-treatment to block the JAK/STAT signaling pathway, which is involved in TNF signaling^44^. We found that treatment with TNF alone elicited very little *IFNL1* expression at similar levels in both parental and ET9 populations, indicating that (a) TNF treatment is insufficient to trigger IFN induction in A549 cells, and (b) that TNF responsiveness does not vary significantly across clonal A549 lines, contrasting with pIC responsiveness. Despite TNF not inducing a large amount of *IFNL1* alone, we observed that TNF pre-treatment synergized with pIC stimulation to drive robust *IFNL1* induction in both cell lines, well beyond what would be expected if the contributions of pIC and TNF were simply additive. Importantly, TNF pretreatment reduced variation in IFN induction between parental and ET9 populations, suggesting that variability in IFN induction between clonal lines can be partially overcome by activating the MAPK signaling network (**Fig. 5A**).

**Figure 5.**
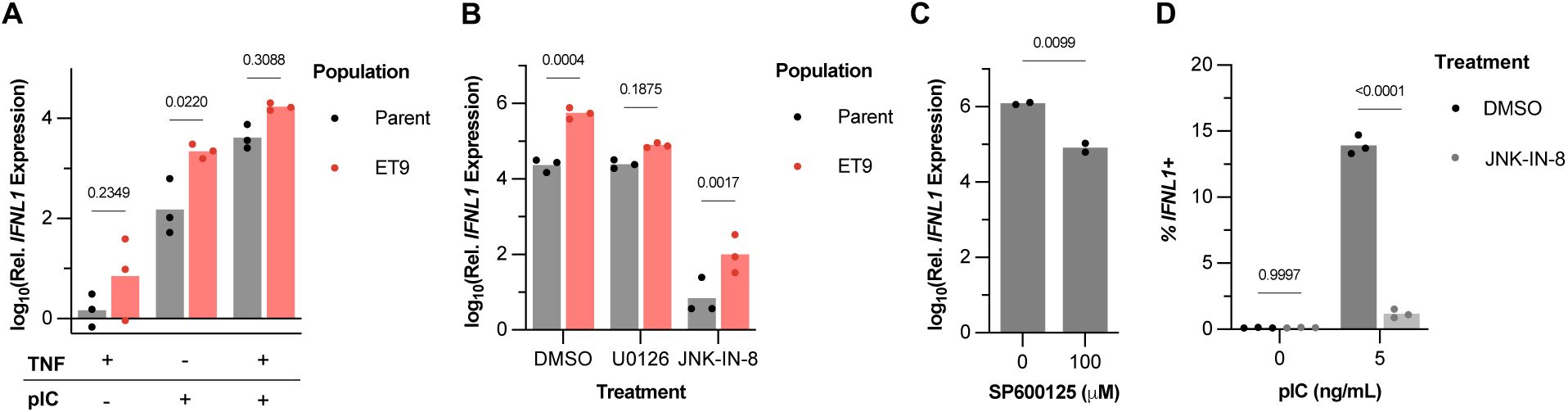
Investigation of MAPK/JNK signaling status effects on IFN expression. **(A)** IFNL1 expression of parental and representative “high” IFNL1 expressor A549 clone in the presence of 10 μM Ruxolitinib with and without 50 ng/mL TNF pretreatment before 0.5 ng/mL pIC treatment for 16 hours. Relative log10(ddCt) fold changes to their ACTB expression and the mean IFNL1 level from the mock of each respective population. Significance determined by Šídák’s multiple comparisons test. **(B)** IFNL1 expression in the parental and representative “high” IFNL1 expressor A549 clone was assessed following pretreatment with 10 μM U0126 or 24 μM JNK-IN-8, followed by 0.5 ng/mL pIC treatment for 16 hours. Relative log10(ddCt) fold changes to their ACTB expression and the mean IFNL1 level from the mock of each respective population. Significance determined by Šídák’s multiple comparisons test. **(C)** IFNL1 expression of A549 cells after 100 μM SP600125 treatment and subsequent stimulation with 5 ng/mL pIC for 16 hours. Relative log10(ddCt) fold changes to their ACTB expression and the mean IFNL1 level from DMSO without pIC. Significance determined by an unpaired t-test. **(D)** HCR-Flow measuring the percentage of cells positive for IFNL1 transcripts after DMSO or 24 μM JNK-IN-8 pre-treatment, then 5 ng/mL pIC stimulation for 16 hours. Significance determined by Šídák’s multiple comparisons test.

We then aimed to inhibit the two MAPKs known to regulate AP-1, JNK and extracellular signal-regulated kinases (ERK), and to examine how this inhibition affected *IFNL1* expression^45,46^. Inhibition of ERK with the MEK inhibitor U0126 did not have a drastic effect on *IFNL1* expression in either population, although it did eliminate the difference between parent and ET9 (**Fig. 5B**). On the other hand, JNK inhibition with the highly selective JNK inhibitor JNK-IN-8^47^ almost completely eliminated *IFNL1* expression in the parent and drastically reduced its expression in ET9 (**Fig. 5B**). We also used another JNK inhibitor, SP600125, to affirm our finding with JNK-IN-8. Although it was less dramatic than with JNK-IN-8 treatment, JNK inhibition using SP600125 did lead to a significant decrease in *IFNL1* expression. (**Fig. 5C**). As a follow-up, we wanted to see if this reduction in expression was from the same number of cells producing less *IFNL1*, or if inhibition of JNK signaling changed the number of cells able to produce *IFNL1* in a population. JNK-IN-8 pre-treatment significantly affected the percentage of cells responsive to pIC stimulation (**Fig. 5D**). Together, these data suggest a role for the MAPK/AP-1 signaling network in governing single-cell heterogeneity in IFN induction potential.

### JNK-IN-8 treatment largely eliminates the transcriptional response to pIC

Given that IFN induction was obliterated by JNK-IN-8 treatment, we next asked whether the entire global transcriptional response to pIC was JNK-IN-8-sensitive. We stimulated A549 cells with pIC in the presence or absence of JNK-IN-8 and the JAK1/JAK2 inhibitor ruxolitinib^43^ (to block effects downstream of IFN signaling) and quantified transcriptional changes using bulk RNA-seq 16 hours after pIC treatment. As expected, ruxolitinib treatment partially decreased pIC-stimulated *IFNL1* expression in A549s, likely due to the role of IFN signaling in enhancing IFN induction (**Fig. 6A**).

**Figure 6.**
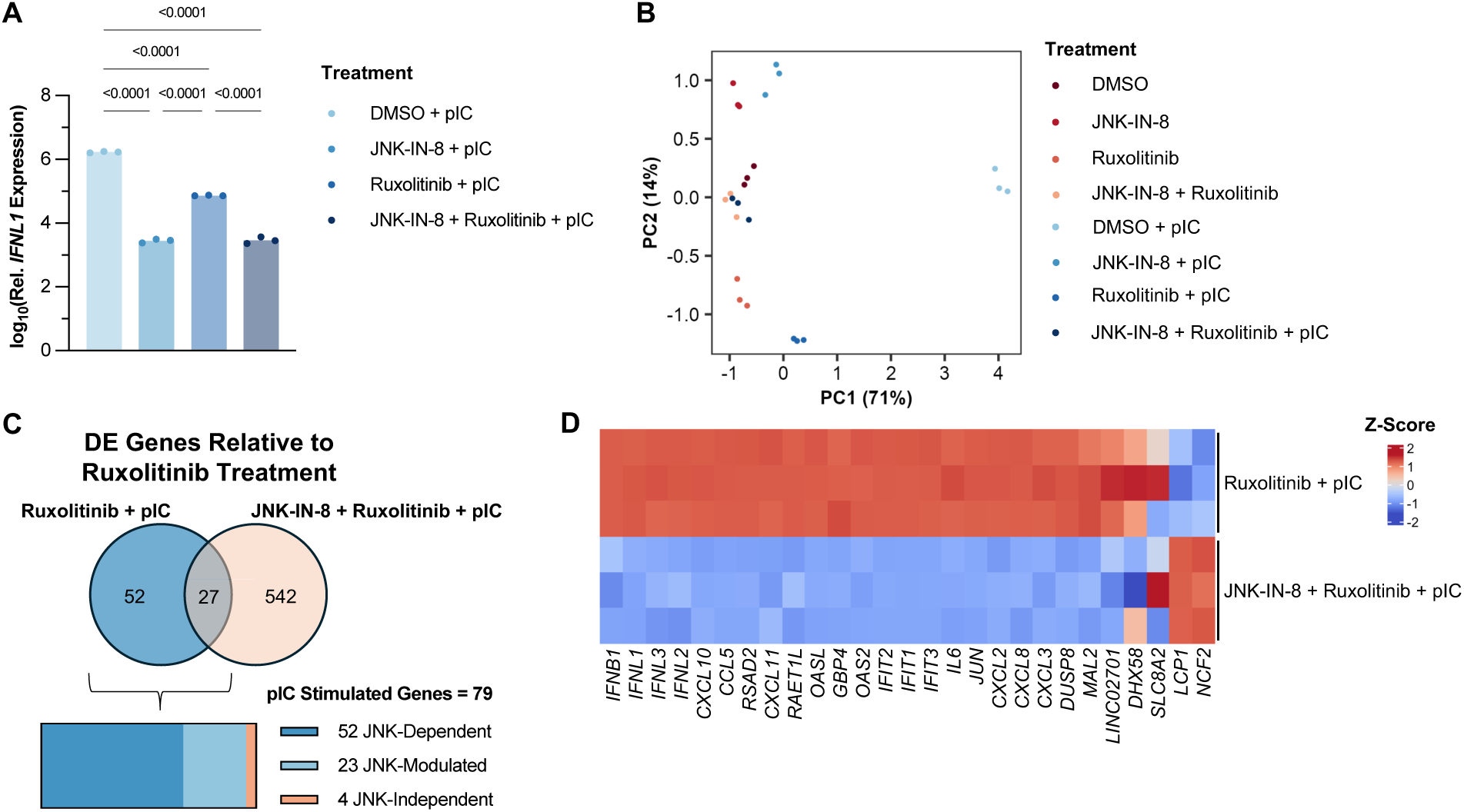
JNK-IN-8 treatment virtually eliminates the innate immune response to pIC. **(A)** IFNL1 expression 16 hours post-treatment with 5 ng/mL pIC. Log10(ddCt) relative to ACTB and DMSO-treated control. Inhibitor treatments with 10 μM ruxolitinib and 24 μM JNK-IN-8 4 hours before pIC stimulation. Significance determined by Tukey’s multiple comparisons test. **(B)** PCA plot comparing the transcriptional profiles of the different treatment groups. **(C)** Venn diagram showing the distinct and shared DE genes in the Ruxolitinib+pIC and JNK-IN-8+Ruxolitinib+pIC treatments. Parts of a whole diagram below detailing what genes of the pIC-stimulated group are JNK-dependent. **(D)** Heatmap comparing the expression of the shared 27 DE genes of Ruxolitinib+pIC and JNK-IN-8+Ruxolitinib+pIC. Gene expression is relative between the two conditions and is represented by the Z-score.

Treatment with either pIC, JNK-IN-8, or ruxolitinib alone had profound, divergent effects on the cellular transcriptome (**Fig. 6B**). Interestingly, JNK-IN-8 and ruxolitinib pre-treatment shifted the transcriptional phenotype of pIC-treated cells to closely resemble that of the DMSO-only controls. This suggested that almost the entirety of the transcriptional response to pIC in A549 cells could be eliminated by inhibiting both JNK-and JAK-dependent signaling.

To specifically define the JNK-dependent portion of the transcriptional response to pIC, we analyzed the differentially expressed (DE) genes in either the ruxolitinib+pIC or JNK-IN-8+ruxolitinib+pIC group relative to the ruxolitinib-only group (**Fig. 6C**). Fifty-two transcripts were significantly upregulated in the ruxolitinib+pIC group but not in the JNK-IN-8+ruxolitinib+pIC group, likely representing the portion of the transcriptional response to pIC that was fully JNK-dependent. Twenty-seven additional transcripts were DE compared to ruxolitinib-only controls in both groups, but only four appeared unaffected by the presence of JNK-IN-8, suggesting that almost the entirety of the transcriptional response to pIC is fully or partially dependent upon JNK signaling (**Fig. 6D**).

## DISCUSSION

In this study, we sought to identify the intrinsic sources of cell-to-cell heterogeneity in IFN induction potential observed across a variety of infection systems^10,12–15,22,23,31^, using the synthetic dsRNA RIG-I ligand pIC. Using multiple complementary approaches, we identified both basal expression of ISGs and the MAPK/AP-1 signaling network as important determinants of single-cell IFN-induction heterogeneity in this system. While we previously identified a role for intrinsic ISG expression (specifically OASL) in influencing IFN induction potential during IAV infection^22^, our data here extend that finding to other stimuli and identify a new role for MAPK/AP-1 signaling in influencing cellular heterogeneity in innate immune function.

Both our longitudinal scRNAseq experiment and our transcriptional analyses of single-cell-derived clones identified the AP-1 transcription factors *JUN* and *FOSB* as correlates of IFN-induction potential. These factors are activated downstream of the MAPK family, an incredibly complex and highly conserved network of signaling molecules that collectively integrate numerous diverse inputs to regulate multiple aspects of cellular behavior, including innate immune function^48–50^. Because of the large number of input signals that can modulate MAPK activity, including cytokines and various sources of cellular stress, the potential involvement of MAPK/AP-1 signaling opens the door to the possibility of a variety of external and internal stimuli contributing to the tuning of IFN induction potential at the single-cell level.

To examine the role of the MAPK network and the downstream AP-1 family transcription factors in governing cellular heterogeneity in IFN induction, we took two approaches, each with significant caveats. First, we tried maximizing MAPK signaling activity by pretreating with TNF in the presence of the JAK/STAT inhibitor ruxolitinib. Our findings confirmed previous work demonstrating that TNF can enhance the production of type I IFNs and their downstream ISGs^44,51–53^. Importantly, they also showed that TNF pretreatment reduced the variability in IFN-induction potential between clonal lines of A549 cells. While this is consistent with a role for MAPK signaling in IFN-induction heterogeneity, the effects of TNF stimulation are many (including activation of IRF1^44,52–54)^, and thus these results do not provide much clarity on the specific players involved.

Second, we tried inhibiting AP-1 function by treating with drugs that target the two main pathways connecting MAPK signaling and AP-1: JNK and ERK. Inhibition of JNK activation nearly eliminated bulk expression of *IFNL1* and nearly every other transcript induced by pIC stimulation. JNK-IN-8 treatment also significantly decreased the percentage of cells capable of expressing *IFNL1* after pIC stimulation. In contrast, ERK inhibition had no appreciable effect on IFN induction, suggesting that the effects of MAPK activity on IFN induction are mediated through JNK. While JNK-IN-8 treatment has been reported to exhibit a high degree of specificity for JNK relative to other kinases^45^, it is impossible to guarantee specificity at the μM concentrations used here. While we found similar effects on IFN induction with the alternative JNK inhibitor SP600125, we cannot rule out that the effects we observed with JNK inhibitors are due to off-target effects. At a minimum, our results point towards a potential therapeutic role for JNK-IN-8 or other JNK inhibitors in the treatment of some interferonopathies.

Our finding that cellular heterogeneity in IFN-induction potential could be stably maintained through multiple generations has important implications for understanding innate immune function within epithelial tissues. If there is significant heterogeneity within populations of epithelial stem cells, this heterogeneity could be passed on through transiently amplifying cells into descendant populations of differentiated epithelial cells^55^. This could potentially result in microenvironments with differing sensitivities to innate immune stimuli across the epithelia.

Though we could not pinpoint a specific mechanism underlying this heritability, it is likely epigenetically encoded. If there were no epigenetic basis for single-cell heterogeneity in pIC responsiveness, we would expect that the numerous generations required to expand individual sorted cells into the clonal lines we examined would eliminate any transient variability in responsiveness seen at the single-cell level. A large body of work has demonstrated that cellular populations can exhibit substantial single-cell heterogeneity in epigenetic states, which gives rise to phenotypic variation^19,21^. These results are also consistent with previous studies demonstrating that IFN expression is subject to epigenetic regulation^20,56–59^ and that single cells isolated from populations during CRISPR knockout generation are often not representative of the population as a whole^60,61^. Our results further emphasize the need for extreme caution when interpreting data generated from single-cell clones (*e.g.,* CRISPR knockouts).

Our results point towards important roles for MAPK/AP-1 signaling and intrinsic ISG expression patterns in determining the fraction of cells capable of responding to viral stimuli by producing IFNs. Because the IFN response is highly non-linear, even small changes in the percentage of IFN-producing cells can have disproportionate effects on the kinetics and magnitude of the downstream antiviral response^62^. The percentage of cells capable of IFN induction effectively sets a threshold of how far viral replication must proceed before triggering an IFN response. Such a system could help avoid unnecessary IFN induction by harmless exposures to PRR ligands, such as might be expected during (a) the limited viral replication that can occur within an immune host, or (b) a transient breakdown in metabolism or containment of host-encoded immunostimulatory RNAs. Future studies should more closely examine how patterns of single-cell heterogeneity in IFN-induction potential vary across different cell populations and environmental contexts.

## LIMITATIONS OF THE STUDY

There are significant limitations surrounding the use of transformed cell lines like the A549 human lung adenocarcinoma cell line used throughout this study. These cells diverge from primary epithelial cells in many important ways, including genomic stability and immune function. Further work is needed both in primary cell systems and *in vivo* to validate and extend the results we report here. As indicated above, it is impossible to rule out off-target effects when evaluating the results obtained using the JNK inhibitor drugs. Both our analyses of single-cell clones and our single-cell pseudotime analysis approaches rely upon correlative results, which do not conclude any causal relationship. More work is needed to firmly establish causal relationships.

## RESOURCE AVAILABILITY

### Lead contact

All additional questions for further information or resources should be directed to the lead contact, Chris Brooke.

### Materials availability

Clonal lines and other reagents are available upon request.

### Data and code availability

All sequencing data can be found at NCBI GEO under accession numbers GSE313066, GSE313067, GSE313068, and GSE313071. The code used for the data analysis in this manuscript can be found at https://github.com/BROOKELAB.

## ACKNOWLEDGEMENTS

We thank the scientists at the sequencing facility in the Roy J. Carver Biotechnology Center for their help with consultation and preparation for our many sequencing experiments. We also thank those at HPCBio, specifically Drs. Chris Fields and Jenny Drnevich, for their guidance in navigating downstream data processing and analysis for the sequencing experiments. Additionally, Dr. Kevin Van Bortle was a great source of advice with the ATAC sequencing analysis. C.B.B was supported by NIH R01 AI139246, NIH R01 AI179910. N.C.W. was funded by NIH R01 AI165475. J.C. was supported by NIH R01 GM089771. S.M. was supported by NSF 2107344.

## AUTHOR CONTRIBUTIONS

C.B. and E.A.T. conceptualized the study. E.A.T., G.S., J.R.C., and J.L., performed all the experiments. E.A.T. and J.S.P. conducted all of the bulk RNA and ATAC sequencing data processing and analysis. E.A.T. processed the single-cell RNA sequencing data for T.M. to perform the temporal-noSpliceVelo analysis, and then E.A.T., with help from J.R.C., analyzed the data further. S.M. and T.M. conceptualized temporal-noSpliceVelo. Q.W.T., J.C., and N.C.W. provided critical reagents. E.A.T. and C.B. wrote the manuscript.

## MATERIALS AND METHODS

### Cells

Human lung epithelial cells (A549) were a gift from Dr. Jonathan Yewdell and maintained in Gibco’s Dulbecco’s Modified Eagle’s Medium (DMEM) high glucose supplemented with GlutaMax and sodium pyruvate, along with the addition of fetal bovine serum (Avantor) for an ending concentration of 8.3%. The cells were cultured at 37°C with 5% CO2.

### Generation of single cell clonal lines

Low-passage A549 cells were collected for FACS and individual cells were distributed into a 96-well flat-bottom plate (Falcon) containing culture media + 5% Penicillin/Streptomycin (Gibco), with 1 cell per well. The wells were then monitored daily until colonies were visible under a microscope. Each well was tracked for whether it contained only one colony or multiple colonies (indicating that the seeding was not perfectly 1 cell per well). Only wells containing a single colony were trypsinized (0.05% Trypsin-EDTA 1X, Gibco) when confluent and expanded into 24-well plates, then into 6-well plates, and finally into T-25 then T-75 flasks. The clones (ET1-11) were numbered based on the order of initial collection.

### Polyinosinic:polycytidylic acid (pIC) treatment

A549 cells were transfected using Lipofectamine 3000 (Invitrogen) with the synthetic double-stranded RNA, pIC (Poly(I:C) LMW, InvivoGen), for the indicated time points. After the specified treatment times, the samples were collected based on the specific downstream quantification method.

### siRNA treatment

We used Dharmacon ON-TARGETplus SMARTpool siRNA (Horizon Discovery) to knock down gene transcripts. The siRNAs used were against human DDX58 (Gene ID: 23586, L-012511-00-0005), TLR3 (Gene ID: 7098, L-007745-00-0005), IFIH1 (Gene ID: 64135, L-013041-00-0005), and non-targeting control (D-001810-10-05). The A549s were transfected with 30 nM siRNA using Lipofectamine RNAiMAX (Invitrogen) according to the manufacturer’s protocol. The cells were incubated for 48 hours, then the RNA was extracted to determine knockdown efficiency, or the cells were treated with pIC as previously described.

### Tumor necrosis factor (TNF) treatment

JAK1/2 inhibition was achieved using ruxolitinib (MedChemExpress), which was first dissolved in DMSO and diluted to 10 μM in cell culture media. The cells were treated with ruxolitinib for 4 hours before TNF, pIC, or both were added to the cells and remained in the media for the entirety of the incubation. Recombinant human TNF (PeproTech) was diluted to 50 ng/mL in culture media, then added to cells 30 minutes before the cells were either collected for western blot analysis or transfected with pIC. The pIC-transfected cells were incubated for 16 hours, after which the RNA was collected for gene expression analysis.

### Drug treatments

All inhibitors were dissolved in DMSO and stored at -70°C in single-use aliquots for long-term storage. Each inhibitor was diluted to its working concentration, with the DMSO concentration not exceeding 1%, in culture media and remained in the culture media throughout the experiment. The cells were incubated with the inhibitors for 4 hours before the addition of their specified stimulus.

### RNA isolation, reverse transcription, and gene expression quantification

The cells were first washed with 1X DPBS (Gibco), then lysed with Buffer RLT (QIAGEN). The samples were then collected and processed using the RNeasy Mini Kit (QIAGEN) according to the manufacturer’s protocol, or frozen at -70°C until further processing could be performed. cDNA synthesis was performed using the Verso cDNA Synthesis Kit (Thermo Scientific) with the provided anchored oligo(dT) primers, according to the user guide. qPCR could then be performed on the QuantiStudio 3 system (Applied Biosystems) with TaqMan reagents (Life Technologies). Gene expression levels were calculated using ΔΔCt, normalizing to ACTB.

### Hybridization chain reaction combined with flow cytometry (HCR-Flow)

HCR-Flow was performed as previously described^29^. As stated, a single cell suspension of A549 cells was prepared by incubating cells with 0.05% Trypsin-EDTA (Gibco) at 37°C for 5 minutes. The cell suspension was washed with DPBS (Gibco) and fixed with 4% formaldehyde (Thermo Scientific) for 30 minutes at 4°C. After incubation, cells were washed twice with PBST and incubated overnight in 70% ethanol. The single cell suspension was washed twice with DPBS + 0.1% Tween20 (PBST) (Thermo Scientific Chemicals) and incubated with amplification buffer (Molecular Instruments) for 30 minutes at 37°C. After incubation, 1 μM of each probe (Molecular Instruments) was added to the cell suspension and incubated overnight at 37°C. Cells were resuspended in probe wash buffer (Molecular Instruments), incubated at 37°C for 10 minutes, and this was repeated three times. Cells were washed once with 5x sodium chloride sodium citrate + 0.1% Tween 20 (SSCT) and incubated in amplification buffer (Molecular Instruments) at RT for 30 mins. After incubation, 3 μM of each snap-cooled hairpin (Molecular Instruments) was added to the cell suspension and incubated at RT overnight. Cells were washed twice with SSCT and analyzed using BD FACSymphony A1.

### Immunofluorescence microscopy and analysis

A549 cells were transfected with 5 ng/mL pIC and incubated for 16 hours before they were washed with DPBS and fixed with 4% formaldehyde (Thermo Scientific) for 30 minutes at 4C. The cells were then washed with PBST three times at RT, permeabilized with DPBS + 0.2% Triton X-100 (Thermo Scientific Chemicals) for 30 minutes, and washed again three times with PBST. Then the cells were blocked with DPBS + 0.1% TritonX-100 + 4% FBS for 30 minutes at RT and incubated with Mouse anti-IRF3 (10949S, CST) in the blocking solution for 1 hour at RT. After washing three times with PBST, the cells were incubated with AlexaFluor 647 AffiniPure Donkey anti-Mouse (715-605-150, Jackson ImmunoResearch) in blocking buffer for 1 hour at RT. The cells were washed three times with PBST, stained with DAPI (Thermo Fisher) in PBST buffer for 10 minutes at RT, then washed three more times in PBST. Finally, the slides were mounted using ProLong Gold antifade reagent (Invitrogen), and a coverslip was set and sealed with nail polish before imaging on a Zeiss LSM 880.

The images were analyzed using Imaris (Oxford Instruments). Briefly, DAPI was used to generate 3D objects, and the mean fluorescence intensity (MFI) of the IRF3 signal within the nuclei was recorded. The MFI of untreated versus pIC-treated cells could then be compared to determine the nuclear translocation of IRF3.

### Bulk RNA sequencing of clonal cell lines

The A549 cell lines ET1-11, along with the parent line, were initially seeded at 1 million cells per well in 6-well plates with three wells per cell line. 24 hours after seeding, the cells were collected, and RNA was isolated as previously described. The Roy J. Carver Biotechnology Center (RJCBC) sequencing facility was responsible for all downstream preparation and sequencing. This included QC, MiSeq (Illumina) titration, cDNA library generation using the TruSeq Stranded mRNAseq Sample Prep kit (Illumina), and sequencing on the NovaSeq 6000 (Illumina) using a S4 flow cell with 2x150nt reads.

### Bulk RNA sequencing of JNK-IN-8, Ruxolitinib, and pIC-treated A549s

The A549 cells were seeded at 1 million cells per well in 6-well plates, with three wells per condition. Cells were either treated with 1% DMSO, 24 μM JNK-IN-8, 10 μM ruxolitinib, or both inhibitors for 4 hours, then 5 ng/mL pIC was added as previously described. The cells were incubated for 16 hours, then collected for sequencing using the same methods. These samples were also processed and sequenced at the RJCBC sequencing facility. The libraries were prepared with the Kapa Hyper Stranded mRNA library kit (Roche) and sequenced using the NovaSeq X Plus (Illumina) on one 10B lane with 1x100nt reads.

### Bulk ATAC sequencing of clonal cell lines

The A549 cell lines ET1-11, along with the parent line, were initially seeded at 1 million cells per well in 6-well plates with three wells per cell line. 24 hours after seeding, the cells were trypsinized and collected on ice for transport to the RJCBC for processing. This includes library preparation with the ATACseq kit from Active Motif and sequencing on the NovaSeq X Plus (Illumina) on one 10B lane with 2x150nt reads.

### Single-cell RNA sequencing of A549s

A549 cells were seeded at 1 million cells per well in 6-well plates. 24 hours after seeding, the cells were treated with 5 ng/mL pIC as previously described. The cells were then collected at their specific time points, transported to the RJCBC on ice, and given to the sequencing facility for further processing and sequencing. The libraries were prepared with the Chromium Next GEM Single Cell Fixed RNA Sample Preparation Kit for Human Transcriptome (10X Genomics) and sequenced using a NovaSeq X Plus (Illumina) on two 10B lanes with 28x150nt reads.

### Modification and use of temporal-noSpliceVelo for the single-cell RNA sequencing dataset

This method of expression-trajectory analysis has been detailed in previous work^22,30^. However, we modified the method to address differences in the samples used. Before estimation of fate probabilities, we removed genes associated with the cell cycle to ensure that we could specifically focus on genes that are not directly associated with the cell cycle. For this, we collected cell-cycle genes^63^ and filtered them. This led to the removal of 38 genes from the 502 genes that were used to train temporal- noSpliceVelo. We could then use the fate probabilities along with the gene expression profiles of each cell for further analysis, which can be found in our code repository https://github.com/BROOKELAB.

